# Reprogramming Adeno-Associated Virus Tropism Via Displayed Peptides Tiling Receptor-Ligands

**DOI:** 10.1101/2022.09.26.509383

**Authors:** Andrew Portell, Kyle M. Ford, Amanda Suhardjo, Joseph Rainaldi, Mark N. Bublik, Milan Sanghvi, Aditya Kumar, Madeleine K. Wing, Nathan D. Palmer, Duy An Le, Nikitha Kalahasti, Amir Dailamy, Prashant Mali

**Affiliations:** Department of Bioengineering, University of California San Diego, USA; Biomedical Sciences Program, University of California San Diego, USA; Department of Chemistry and Biochemistry, University of California San Diego, USA; School of Biological Sciences, University of California San Diego, USA

## Abstract

Adeno-associated viruses (AAVs) are common gene therapy vectors, however, their effectiveness is hindered by poor target tissue transduction and off-target delivery. Hypothesizing that naturally occurring receptor-ligand interactions could be repurposed to engineer tropism, we fragmented all annotated protein ligands known to bind human receptors into tiling 20-mer peptides and displayed these onto the surface loops of AAV5 and AAV9 capsids at two sites. The resulting four capsid libraries, comprising >1 million AAV variants, were screened across 9 tissues in C57BL/6 mice. Tracking variant abundance, we identified >250,000 variants which packaged into capsids, and >15,000 variants which efficiently transduced at least one mouse organ. We individually validated 21 AAV variants with 74.3% of the organ tropism predictions accurately reproducing, confirming overall screen efficacy. Systematic ligand tiling enabled prediction of putative AAV-receptor interactions, which we successfully validated by targeted genetic perturbations. Comprehensive peptide tiling also enabled examination of homologous peptide activity. Interestingly, we observed functional peptides tended to be derived from specific domains on ligands. Notably, certain peptides also displayed consistent activity across mice strains, capsid insertion contexts, and capsid serotypes, including novel immune orthogonal serotypes. Further analyses of displayed peptides revealed that biophysical attributes were highly predictive of AAV variant packaging, and there was a machine learnable relationship between peptide sequence and tissue tropism. We anticipate this comprehensive ligand peptide tiling and display approach will enable engineering of tropism across diverse viral, viral-like, and non-viral delivery platforms, and shed light into basic receptor-ligand biology.

## INTRODUCTION

Nucleic acid and protein based therapeutics to modulate healthy and diseased states are poised to enable the next frontier of human medicine. However their successful deployment is contingent on our ability to deliver them efficiently and in a safe and targeted manner (*1*). In this regard, a host of viral and non-viral delivery formulations have been developed (*2–5*), but the ability to programmably modulate their tropism remains challenging (*6, 7*).

Here we focus on adeno-associated viruses (AAVs) which have emerged as a leading vector for gene delivery in clinical applications (*8–10*). While multiple AAV-mediated therapies have achieved regulatory approval (*11*), efficient directing of treatment to target tissues is challenging with systemic injection. To overcome this, high viral titers are often used in treatments, which has in turn been associated with potential for hepatotoxicity in clinical trials (*12*). Localized injections are also problematic, often requiring invasive procedures with the potential for organ damage and long recovery times. Due to these delivery challenges, some gene therapeutics have elected to, where feasible, pursue *ex vivo* treatment designs to overcome targeting issues, but this in turn can lead to dependency on complex lab procedures and high manufacturing costs (*13*).

There are thus numerous ongoing research efforts to improve *in vivo* therapeutic targeting. In this regard, groups in the field have progressively engineered AAV variants to specifically target tissues such as the brain (*14, 15*) and muscle (*16–18*). This has predominantly been accomplished using strategies of iteratively screening random peptides inserted into the AAV capsid (*19–21*), capsid shuffling (*22–26*), randomly mutagenizing the capsid sequence as a whole (*27–30*), or direct chemical engineering (*31, 32*). Although mutagenizing AAV capsids via random oligomers has yielded functional capsids with exciting and novel properties, a stochastic mutational screening strategy limits our ability to predict future functional variants, and thus rational and programmable engineering of viral phenotypes remains an elusive goal.

Towards rational engineering of viral function, deep mutational libraries and associated screens of function have enabled systematic mapping of capsid mutation fitness (*33, 34*), providing critical information which can be used to predict future variant activity. Additionally, defined libraries of pooled oligonucleotides have been used to insert gene fragments derived from proteins with known affinity to synapses into the AAV capsid (*35*), with the goal of improving retrograde axonal transport. While these methodologies have provided important insights for AAV engineering, much is still unknown regarding how AAV genotype impacts packaging and tissue transduction. Consequently, there is a critical need for systematic datasets mapping AAV genotype to clinically relevant properties such as organ specificity. Given the clinical danger of hepatotoxicity (*12*) and other efficacy issues related to off-target transduction (*36*), leveraging screening technologies to yield highly specific AAV capsids has great value to the medical and scientific community.

With a goal to develop a rational and potentially programmable strategy for enabling tissue targeting, we hypothesized that receptor-ligand interactions which mediate the spectrum of naturally occurring cell-protein and cell-cell interactions could be repurposed for engineering AAV tropism. However, as neither the interaction interfaces nor the cell-type specificity profiles of receptor-ligand interactions are fully mapped, to comprehensively interrogate as well as engineer these, we thus leveraged our recently developed *PepTile* approach for mapping and targeting putative protein interaction domains (*37*). Specifically, we fragmented all annotated protein ligands known to bind human receptors into tiling short peptides and displayed these onto surface loops of AAV capsids, an approach we term *AAV-PepTile*. Short peptides grafted into stabilizing molecular scaffolds can recapitulate local protein domain structure (*38*), and as protein-protein interface sites are typically 1200-2000 Å^2^ (*39*), with specific peptide hot loops ranging from 4-8 amino acids (AAs) contributing maximally to the binding energy of protein-protein interactions (*40*), we hypothesized that 20 amino acid peptide insertions into the AAV capsid could drastically alter its binding ability, and therefore, its transduction profile *in vivo.* While insertion of still longer peptides could in principle more faithfully recapitulate local ligand structures (*37, 41*), AAVs do not generally tolerate larger than 20-25 amino acid insertions without severely compromising capsid packaging efficiency (*42*). Based on this, we proceeded with a 20AA peptide + flanking 2AA linker insertions, all together constructing a library of >1 million AAV variants by inserting corresponding oligonucleotide pool synthesized gene fragments coding for potential receptor-ligands and cell membrane permeable proteins into one of two surface loops on AAV5 and AAV9. Unlike random peptide libraries, ligand tiling enabled robust quantitation of tissue transduction rates for all variants screened. Furthermore, systematic examination of the activity of similar peptides generated predictions of putative receptor interactions driving AAV variant tropism. Quantifying transduction rates across nine organs, we identified extremely specific variants targeting the brain and lung, as well as muscle and heart targeting variants with broader organ transduction. The resulting data linking AAV variant genotype to packaging efficacy and tissue specificity expands our understanding of the AAV fitness landscape and provides a unique resource from which further data-driven engineering efforts could be built.

## RESULTS

### A systematic library of AAV variants displaying fragmented proteins

To create our libraries we chose AAV5 and AAV9 as the starting serotypes. This was due to their established clinical utility, as well as two key characteristics: one, AAV5 is more evolutionarily distant to other AAV serotypes in clinical use, and has previously been shown by us to be immune orthogonal to other prevalent AAV serotypes (thereby enabling their sequential re-dosing) (*43*); and two, AAV9 has been used extensively for clinical trials and has been shown to cross the blood-brain barrier, outperforming other AAV serotypes in most tissues (*11, 44–47*). To generate our library of diverse AAV variants, we synthesized a DNA oligonucleotide pool of 275,298 gene fragments (**Fig. 1a-b**, **Methods**). Each gene fragment coded for a 20 amino acid peptide derived from the coding sequence of ligands with known extracellular receptors, or a gene predicted to have cell-penetrating or internalizing properties (**Fig. 1a-b, Methods**). Protein ligands were sourced from the Guide to Pharmacology database, an expertly curated list of pharmacological targets and their associated ligands (*48*), and cell-penetrating/internalizing functionality was inferred through text mining of UniProt entries. Examples of protein classes identified as having potential internalizing function included toxins (*49*), histones (*50*), granzymes (*51*), viral receptor binding domains (*35*), and nuclear localization signal (NLS) domains (*52*). Mouse and human genomes share 80% of their protein coding genes (*53*), with 85% amino acid sequence identity between orthologs (*54*). These genomic similarities extend to the receptors for ligands in our libraries (e.g. 86% of mouse GPCR proteins have human orthologs (*55*)), implying many human ligands will be similarly functional across human and mice contexts. After synthesizing the pool of single-stranded oligonucleotides coding for these gene fragments, it was amplified to double-stranded DNA via PCR, digested, and ligated into two distinct locations on the AAV5 and AAV9 *cap* genes (**Methods**). Seamless cloning was enabled by type IIS restriction enzymes cutting outside their recognition sequence. PaqCI sites engineered to yield compatible overhangs (coding for glycine-serine 2 amino acid linkers) were inserted at the ends of the peptide coding DNA library and on the AAV5/AAV9 *cap* plasmid DNA at sites coding for two distinct surface loops, hereon referred to as loop 1 and loop 2 **(Supplementary Fig. 1a)**. Surface loop 1 (AA 443 and 456 on AAV5 and AAV9, respectively) and loop 2 (AA 576 and 587 on AAV5 and AAV9, respectively) were chosen as peptide insertion locations due to their distance from the viral particle core facilitating potential receptor engagement (**Fig. 1b**). While many previous AAV engineering efforts have focused on inserting peptides at surface loop 2 (*16, 56*), loop 1 sites have been shown to accommodate large insertions including even a full length fluorescent protein (*57*). Collectively, we produced four libraries of variants, spanning two AAV capsids, with two loop insertion sites each. In addition to protein coding gene fragments, we also included 444 stop codon containing gene fragments as negative controls. We utilized this defined oligonucleotide library synthesis methodology to enable quantitative inference of variant packaging and transduction efficiencies. The starting plasmid libraries were sequenced to establish initial relative variant abundances, and packaging efficiencies were quantified via comparison to this initial baseline (**Methods**).

**Figure 1.**
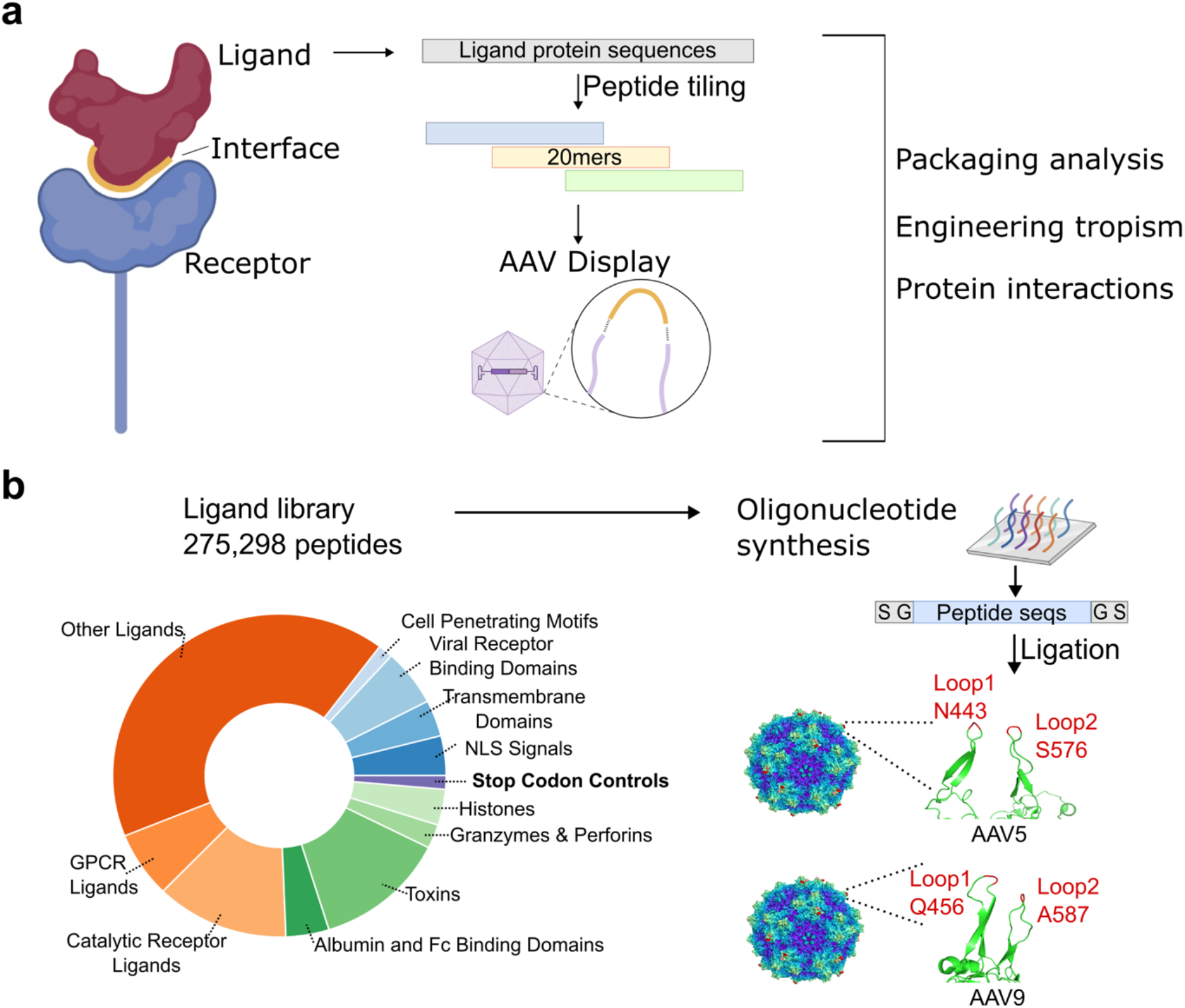
Design of AAV libraries displaying peptides tiling receptor-ligands. **(a)** Schematic of approach for rationally engineering and characterizing AAV variants. Ligand protein sequences derived from all annotated receptor-interacting ligands were systematically tiled into 20 amino acid peptides and inserted into surface-exposed loops of AAV capsids. These engineered variants were then assessed for their packaging capacity*, in vivo* tropism, and enhanced protein interactions. **(b)** Protein class distribution of the ligands which were tiled to compose our screening library. These include known receptor-interacting ligands (orange), cell membrane permeable proteins (green), protein domains (blue), and stop codon containing negative controls (purple). Peptide sequences were generated via pooled oligonucleotide synthesis and inserted into 4 distinct loop regions: AAV5-Loop1 (N443), AAV5-Loop2 (S576), AAV9-Loop1 (Q456), and AAV9-Loop2 (A587) to generate over 1 million AAV variants. Capsid surface residues are colored according to their distance from the core of the capsid with the insertion site flanking residues shown in red.

### Biophysical drivers of AAV capsid formation

To quantify how well different AAV *cap* variants package into functional capsids, we generated recombinant AAV particles with our engineered AAV5 and AAV9 *cap* plasmid libraries via transient triple transfection of HEK293T cells (**Fig. 2a**, **Methods**). These viral particles were treated with benzonase to degrade residual plasmid DNA, and then subjected to next generation sequencing (NGS) to quantify relative variant abundance. Packaging efficiency was quantified by ranking AAV variants by the log_2_ fold change (log_2_FC) of their relative capsid abundance compared to their count in the plasmid pool (**Fig. 2b, Methods)**. Utilizing this method, we identified over 250,000 AAV variants which package efficiently into functional AAV capsids with a log_2_FC > 0. To validate this packaging metric quantified from screening data, we individually produced 25 AAV capsids including 23 identified as successful packagers (log_2_FC >0) and 2 identified as non-packagers (log_2_FC < 0). In these individual validations, the 2 non-packagers yielded >10-fold lower titer than those identified as packagers, thus providing confidence in our AAV packaging metric. Consistent with their disruption of the AAV capsid structure there was also a depletion of non-functional stop codon control AAV variants in the capsid pool, and importantly this confirmed lack of library cross-packaging during AAV production **(Supplementary Fig. 1b)**.

**Figure 2.**
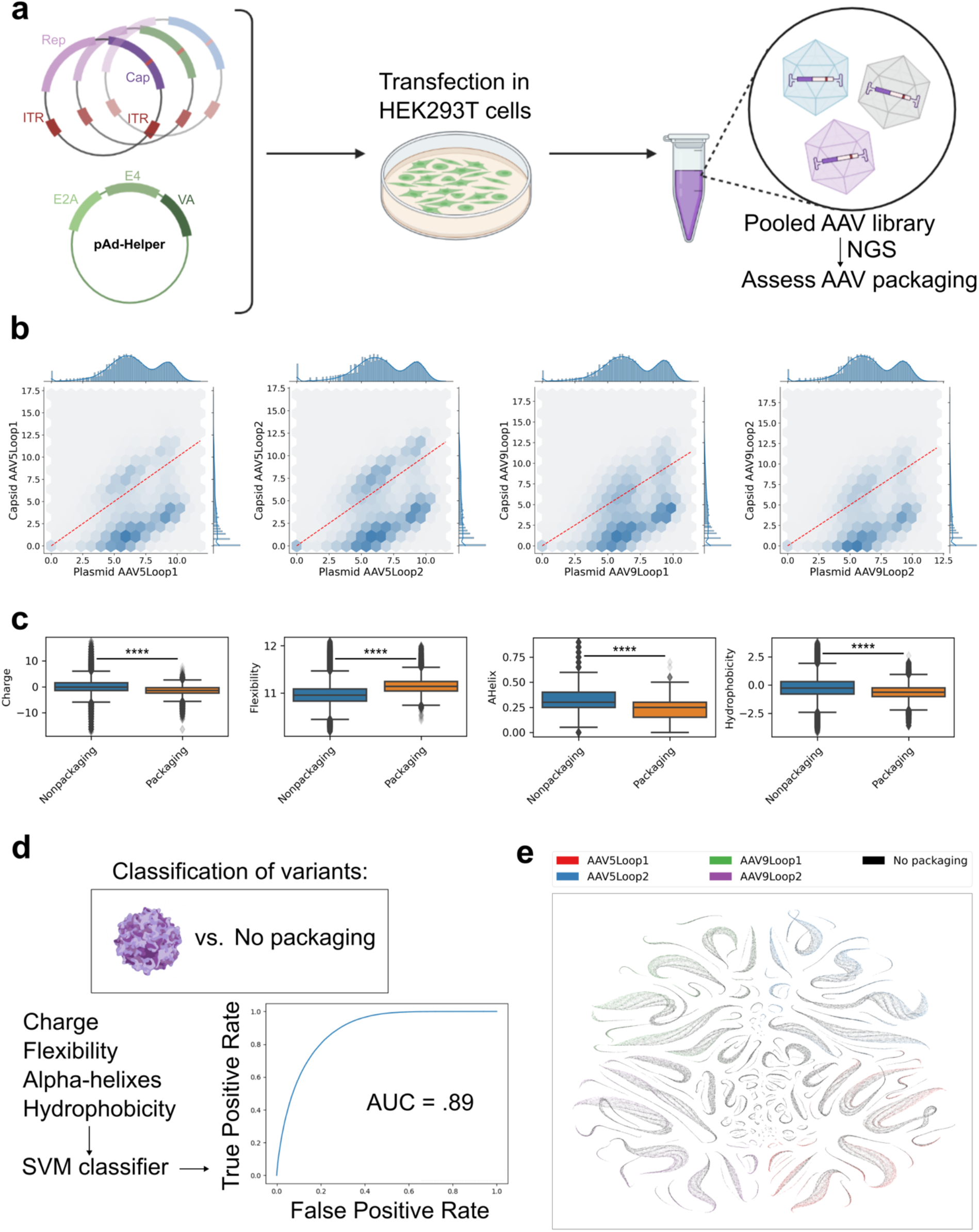
AAV library packaging analyses reveal biophysical features contributing to capsid fitness. **(a)** Schematic illustrating recombinant production of pooled AAV libraries in HEK293T cells. **(b)** Normalized abundance for each inserted peptide in the plasmid libraries versus DNA isolated from recombinantly produced AAV variant capsid libraries. Dotted-red line shows where the plasmid abundance is equal to the capsid abundance. **(c)** Distribution of peptide biophysical parameters relevant to packaging. Peptide charge, alpha-helical content, flexibility, and hydrophobicity distributions are shown for peptides which are enriched in the capsid pool (“Packaging”) and those which are depleted (“Non-Packaging”) **(Methods)**. Statistical significance between groups was calculated via a T-test (****p<0.0001). **(d)** Biophysical parameters of inserted peptides were used as features to train a support vector machine (SVM) classifier predicting which AAV variants successfully package into capsids. The receiver operating characteristic curve is shown for the resulting model, with an area under the curve of 0.89. **(e)** UMAP embedding for each AAV variant, colored by packaging status. Inserted peptide charge, alpha-helical content, flexibility, and hydrophobicity were used as input features for the embedding.

Next, to better understand what features drove successful capsid formation, we examined the biophysical characteristics of the inserted peptides that yielded AAV variants that packaged successfully. We found that peptide charge, flexibility, alpha-helical content, and hydrophobicity were all significantly different in packaging AAV variants versus AAV variants unable to package (**Fig. 2c**). The set of successfully packaged variants had a narrower charge distribution than the variants unable to package, suggesting peptides with extreme charge densities have a negative impact on capsid formation. Successfully packaging AAV variants also had inserted peptides with higher flexibility, lower alpha-helical content, and lower hydrophobicity than the variants unable to package. The observed depletion of hydrophobic peptide displaying variants is consistent with the solvent exposed nature of the AAV surface loops.

To build an integrated model predicting if AAV variants will package based on the biophysical features of the inserted peptides, we next trained a support vector machine classifier (*58*) using the charge, flexibility, alpha helical content, and hydrophobicity of the peptides in our dataset (**Fig. 2d, Methods**). While all of these biophysical features were significantly different when comparing packaging versus non-packaging AAV variants, the magnitude of this difference was relatively modest for each individual feature (**Fig. 2c**). However, collectively these features were sufficient to train a model which could differentiate between packaging and non-packaging AAV variants (area under the receiver operating characteristic curve = 0.89, **Methods**). For each AAV variant, embedding the inserted peptide’s charge, flexibility, alpha helix content, and hydrophobicity into two dimensions using uniform manifold approximation and projection (UMAP) (*59*) enabled visualization of this class separability (**Fig. 2e**). The resulting embedding laid out AAV variants into distinct clusters, indicating that while each underlying biophysical feature is continuous, there are separable groups of AAV variants with similar biophysical features. AAV variants which package tended to cluster with other packaging variants in this unsupervised embedding, further supporting the predictive power of these four biophysical features. This thorough quantification and analytical framework for assessing AAV packaging, which is enabled by the diversity and length of our inserted peptide library, is of critical translational importance for identified AAV variants as AAV production costs are directly related to their packaging capability.

### High-throughput mapping of engineered AAV tissue tropism

Having produced libraries of recombinant AAV particles packaging their own *cap* genes, we next injected these four viral pools into C57BL/6 mice in duplicate (**Fig. 3a**). After two weeks, mice were sacrificed and the peptide-containing region of the AAV *cap* gene was amplified from DNA isolated from mouse liver, kidney, spleen, brain, lung, heart, skeletal muscle, intestine, and pancreas. We quantified the relative abundance of each AAV variant via NGS, and calculated the log_2_ enrichment of each variant relative to its abundance in the capsid pool, observing good correlation between replicates (**Supplementary Fig. 1c**). Using this data, we identified over 15,000 variants which efficiently infected at least one mouse tissue (**Fig. 3b, Methods**). The spleen and liver were the most frequent tissue targets of the infectious variants, consistent with the established wild-type (WT) tropism of AAV5 and AAV9 towards the liver (*60*), as well as more recent research showing AAV5 and AAV9 readily transduce the spleen (*61*). We observed the fewest AAV variants targeting the skeletal muscle and brain, in line with the high therapeutic AAV doses needed to achieve clinical efficacy for muscle targeting gene therapies (*62*), and the challenge of delivery across the blood-brain barrier (*63*). Finally, we also made the interesting observation that a substantial number of our identified AAV variants were comprised of the same peptide inserted across different AAV capsids and insertion sites, giving credence to our hypothesis that tropism reprogramming is, at least partially, peptide-specific (**Fig. 3b**).

**Figure 3.**
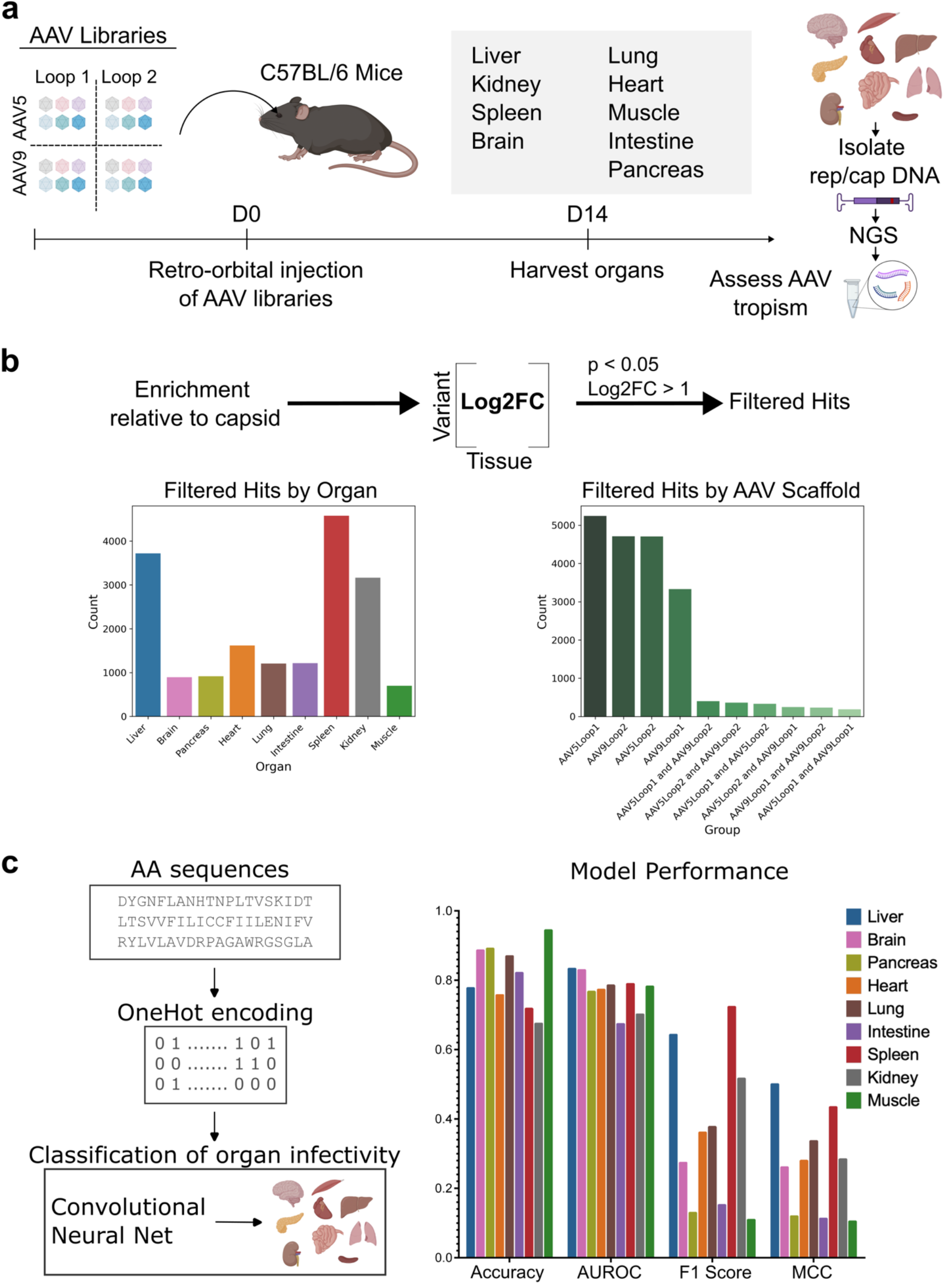
*In vivo* screen analyses enable predictive computational models of tropism. **(a)** Overview of *in vivo* screening methodology. The four AAV variant libraries were injected retro-orbitally into C57/BL6 mice in duplicate. Two weeks post-injection, nine organs were harvested from each mouse and the inserted peptide-containing region of the AAV capsid was amplified and subjected to next generation sequencing. **(b)** Overview of screen results. Significantly enriched AAV variants in a particular organ were defined as those with a log_2_FC > 1 and an FDR-adjusted p-value < 0.05. Bar plots show the number of significantly enriched variants detected per organ, as well as a comparison of peptide hits for each loop insertion site, for both AAV5 and AAV9. **(c)** Overview of classification model predicting AAV tissue tropism from peptide sequence alone. Inserted peptide sequences were converted to a binary one-hot encoding (for each peptide, 20 rows corresponding to position, and 20 columns corresponding to presence of a particular amino acid). This one-hot encoding scheme was then used as input to a convolutional neural network (CNN) to predict organ targeting. Model performance was separately evaluated on each organ, via accuracy, area under the receiver operator characteristic curve (AUROC), F1 score, and Matthews Correlation Coefficient (MCC). Models were trained on ⅔ of the data, and the remaining ⅓ was held out as a validation dataset to evaluate performance.

Towards mapping and understanding the tissue-transduction patterns mediated by the inserted peptides, we examined the feasibility of training predictive models linking inserted peptide sequence to tissue tropism. Inspired by contemporary work using convolutional neural networks (CNN) to predict antibody specificity (*64*), we trained a CNN multi-label classifier to predict AAV tissue tropism using one-hot encoded inserted peptide sequences as input features (**Fig. 3c, Methods**). To evaluate performance, the model was trained using a random selection of ⅔ of the significantly enriched AAV variants, and evaluated on the ⅓ testing dataset. This CNN model architecture had good performance across all organs, with a minimum accuracy of 72% in the kidney (**Fig. 3c**). We observed the highest F1 scores and Matthews Correlation Coefficients (MCC) for the liver and spleen (**Fig. 3c**), likely due to the high number of liver and spleen targeting variants we identified in the pooled screen (**Fig. 3b**). We observed that for the liver and spleen the CNN had only modest reduction in F1 scores when evaluated on peptides very different (greater than 10 edits away) from any training examples, implying the models had learned generalizable features relevant to AAV transduction. For other organs, the models were much less accurate when predicting the tissue tropism of peptides very different from the training data, likely due to the reduced amount of positive training examples. Collectively, the ability to predict AAV tropism supports the conclusion that the inserted peptides are mediating retargeting of tissue-tropism, and that there is a learnable relationship between peptide sequence and tissue tropism.

### Engineered AAV variants with clinically relevant tissue tropism

Like the WT scaffolds from which they were derived, transduction of multiple organs is a near ubiquitous phenotype among the infectious AAV variants identified (**Fig. 4a**). Significant transduction of the liver and spleen was observed for the majority of infectious variants regardless of which other organs were co-transduced. This is true even for variants with insertions in surface loops known to be involved in WT capsid receptor binding (*65, 66*). While liver and spleen targeting were near ubiquitous, we were able to identify variants which specifically target the liver/spleen plus one other organ, as well as variants which transduced all tissues at high levels (**Fig. 4a**). When variants were hierarchically clustered based on their tissue detection levels, we observed that variants derived from the same sub-library tended to cluster together, suggesting that the tissue specificity of the wild-type scaffold was at least partially a determinant of engineered variant tropism. Hierarchical clustering of the organ samples resulted in replicates clustering together, giving further confidence to the reliability of the screen results. To visualize the overall screen results, we embedded the tissue detection levels for each variant into two dimensions using UMAP, coloring the variants by the organ they most readily transduce (**Fig. 4b, Methods**). In this reduced dimensional space, organ specific clusters can be readily identified, with the liver and spleen targeting variants especially prominent.

**Figure 4.**
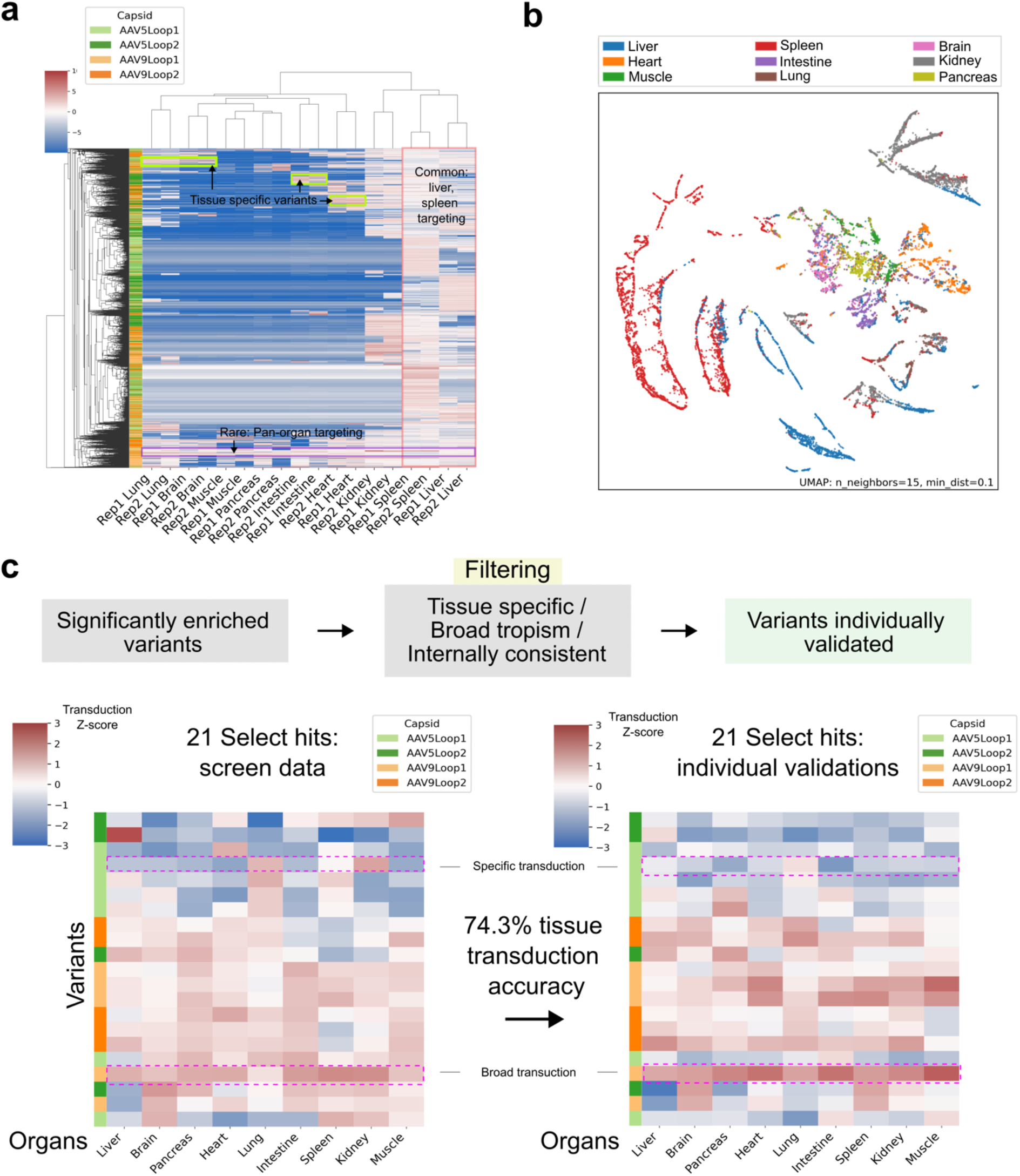
*In vivo* screen identified AAV variants demonstrate reprogrammed tropism. **(a)** Heatmap showing log_2_FC values for each AAV variant which was significantly enriched in at least one organ. Rows are individual variants, and columns are organs (n=2 per organ). **(b)** UMAP embedding of significantly enriched AAV variants. Each dot represents a variant, colored by the organ with max log_2_FC. **(c)** AAV capsids were chosen for validation from the pool of significant hits on the basis of their tissue specificity, broad tropism, and/or internal consistency. Internal consistency was quantified by counting the number of similar (>50% homology) inserted peptides also detected as hits for a given organ (**Methods**). AAV variants were characterized structurally via transmission electron microscopy, and functionally via delivery of the mCherry transgene *in vivo.* Heatmaps depict all variants chosen for validation (n=21), with the left heatmap showing the Z-normalized log_2_FC values from the pooled *in vivo* screen (n=2), and the right heatmap showing the Z-normalized mCherry expression quantified by RT-qPCR, relative to AAV9 (n=2) (**Methods**).

To confirm the tissue tropism of the novel AAV variants identified via the pooled screen, we next individually produced and validated 21 variants by quantifying their ability to package and deliver an mCherry transgene *in vivo* (**Fig. 4c**). All 21 variants were significantly enriched in at least one organ, and we prioritized choosing variants for validation which were internally consistent within the screening data: consistent AAV capsids being defined as hits where we identified other variants with similar inserted peptides enriching in the same organ (**Methods**). Variants were initially characterized by *in vivo* quantification of tissue tropism at the mRNA level (via RT-qPCR quantification of mCherry transgene expression) (**Fig. 4c**). The tissue tropism of the variants largely recapitulated the screen predictions (**Fig. 4c**), with 74.3% of our tissue tropism predictions matching expectations (**Methods**). We identified AAV variants which specifically targeted hard to infect organs such as the muscle, lung, and brain, while simultaneously de-targeting away from the liver (**Fig. 4c**). Notably, protein level quantification of mCherry delivery to the liver, quantified via fluorescent microscopy, confirmed excellent concordance between mRNA and protein measures of tissue transduction (R^2^=0.97, **Supplementary Fig. 2a**). Altogether, across the variants individually tested, we found 9/21 variants exceeded AAV9 infectivity in at least one organ. We identified variants which exceed AAV9 infectivity in all organs except the liver and pancreas, which had max relative transduction of 98.8% of WT AAV9 and 82.2% of WT AAV9, respectively. Additionally, we found that 18/21 variants had less than half the liver transduction of WT AAV9, with three variants below 5% AAV9 liver transduction levels indicating clinically important liver de-targeting. Finally, to increase confidence in the fidelity of our individually validated variants in C57BL/6 mice, we also individually validated 3 of the above 21 variants in BALB/c mice. Notably, their relative tropism was consistent across the two mouse strains **(Supplementary Fig. 2b)**.

### Mechanistic insights into AAV reprogramming

Having confirmed the efficacy of our AAV variants, we next explored the mechanisms underlying their reprogrammed tropism, and in particular, investigated whether our hypothesis that AAV variants displaying peptides derived from distinct ligand domains could indeed drive their transduction patterns. First, we sought to interrogate how peptides derived from specific ligand protein regions alter AAV tropism. This was motivated by our previous observation that tiled peptide enrichment patterns in a screen can yield insight into functional domains of the proteins from which they were derived (*37*). To do this, we interrogated tissue-specific AAV variants identified in the screen, focusing first on a lung-specific variant (termed as AAV9.DKK1) that was derived via display of a peptide fragment from the *DKK1* gene on the AAV9-Loop2 scaffold. Specifically, as shown in **Fig. 5a** **(top left)**, we observed significantly enhanced lung transduction from peptides derived from a specific domain in the N-terminus of the protein. Upon confirming that this variant is packaged into a functional capsid via electron microscopy **(Fig. 5a, top right)**, we quantified its transduction profile *in vivo*. This confirmed that AAV9.DKK1 was indeed an efficient lung transducing variant with greater than 2-3 fold higher expression than AAV9, and with consistent de-targeting across all other organs compared to AAV9 when quantified at both the RNA and protein level **(Fig. 5a, bottom left and right)**. Next, we investigated whether the DKK1 displayed peptide could lend insight into the mechanism of transduction. Interestingly, we observed that the N-termini region of lung enrichment **(Fig. 5a, top left)** was centered on an evolutionarily conserved, linear peptide motif known to mediate binding to the low density lipoprotein receptor-related proteins 5 and 6 (LRP5/6) (*67*). As HEK293T cells robustly express LRP6, we thus utilized lentiviral-mediated CRISPR-Cas9 to disrupt LRP6 expression in these cells to investigate the relative transducability of the AAV9.DKK1 variant in this system. **(Fig. 5b, Methods)**. Specifically, cells transduced with either non-targeting control (NTC) guides, or *LRP6* targeting guides were subsequently transduced with either AAV9.DKK1 or wild-type AAV9. Using flow cytometry to quantify AAV delivery of the mCherry transgene, we observed a significant reduction in the infectivity of AAV9.DKK1 in the LRP6 knockout population, with no corresponding reduction in the infectivity of wild-type AAV9 (**Fig. 5b**). The LRP6-dependent *in vitro* infectivity of AAV9.DKK1, combined with the known interaction between LRP6 and the N-termini of DKK1 (see crystal structure, **Fig. 5b**), and its known robust expression in lung alveolar cells (*68*), together gives strong support to our hypothesis that LRP6 mediates, at least in part, the reprogrammed tropism of AAV9.DKK1.

**Figure 5:**
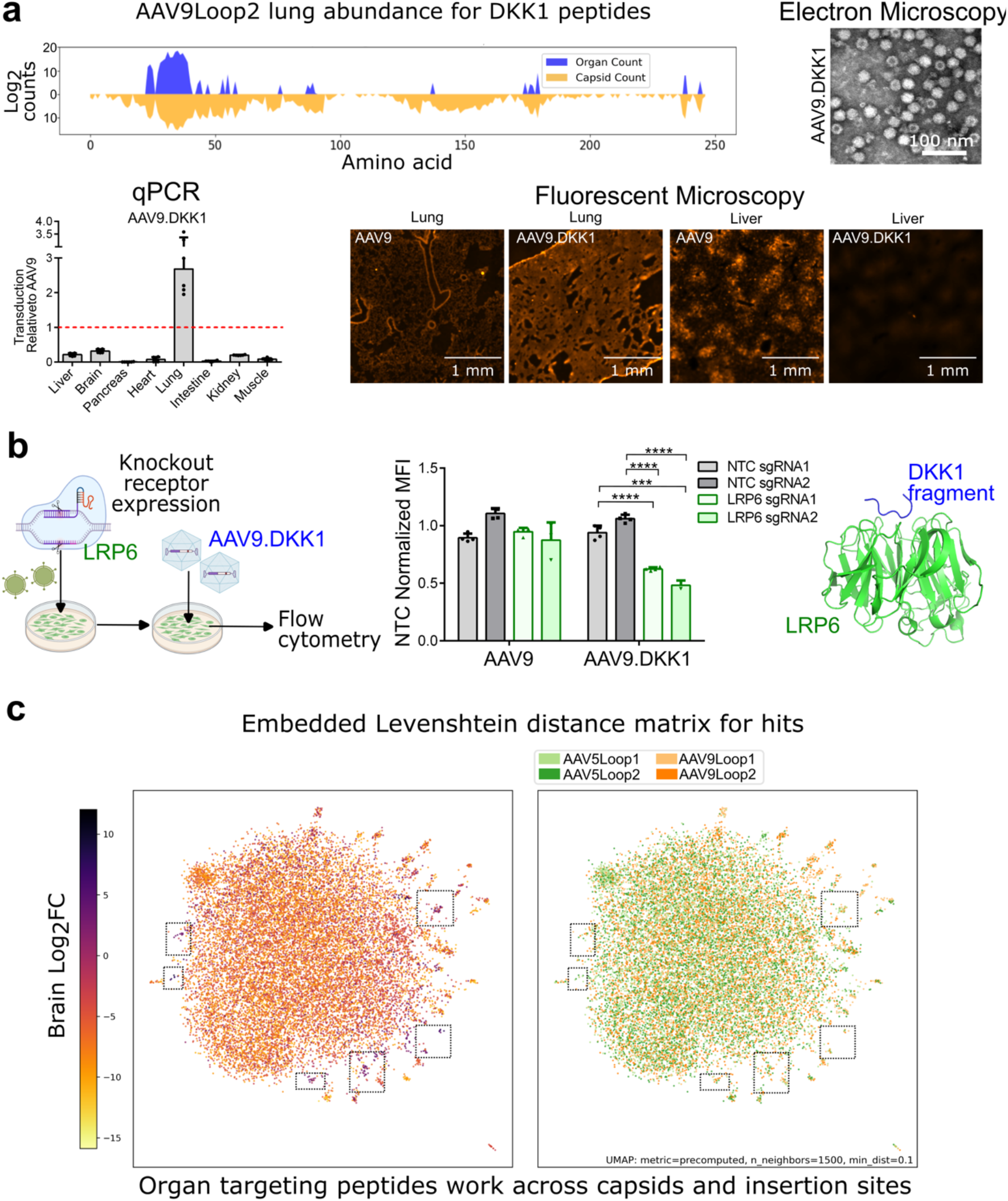
Characterization and mechanistic exploration of AAV variants with displayed ligand peptides. **(a)** Full characterization experiments for the variant AAV9.DKK1. In the top left, lung screen count values for all AAV9Loop2 variants with inserted DKK1 derived peptides are shown. The x-axis indicates the position in the DKK1 structure a given peptide starts on. Shown in blue are the lung counts, and shown in orange are the capsid counts. Bar plot shows the RT-qPCR quantification (n=2) of mCherry transgene expression via individual validation, normalized to that of AAV9. Also shown are electron microscope images for AAV variant capsids, as well as fluorescent microscopy of mCherry protein expression levels in the lung and liver. **(b)** Knockout experiments to validate AAV9.DKK1 receptor dependency. Cas9-containing lentivirus was produced with either non-targeting controls (NTC) and LRP6 targeting sgRNA (n=2 each). HEK293T cells were then transduced with lentivirus and positively selected with puromycin. Lentivirally transduced cells were then plated in a 24-well plate and transduced with either wild-type AAV9 (4×10^9^ viral genomes) or AAV9.DKK1 virus (1×10^9^ viral genomes). Flow cytometry was then used to quantify the mean fluorescence intensity (MFI) of the infected cells. Shown to the right is the crystal structure of a 7-mer peptide DKK1 peptide (contained within AAV9.DKK1) in complex with LRP6 (*67*). Statistical significance between groups was calculated via a t-test (*p<0.05, **p<0.01, ***p<0.001, ****p<0.0001). **(c)** Significantly enriched AAV variants from all capsids were projected into 2 dimensions via UMAP. The distance between points is calculated explicitly via the Levenshtein distance of the amino acid sequences of the inserted peptides, and subsequently embedded via UMAP (**Methods**). AAV variants are colored by their log_2_ fold change in the brain. Select variant clusters which are highly enriched in the brain are highlighted. AAV variants are colored by capsid insertion site in the embedding on the right.

Building on these observations, and focusing on the brain as an exemplar, we next sought to examine more broadly the extent to which the inserted peptides were mediating re-targeting. First, we constructed a distance matrix quantifying the similarities between all peptide hits identified as significantly enriched in at least one organ (**Fig. 5c, Methods**). By projecting this distance matrix into two dimensions, we were able to visualize distinct clusters of homologous inserted peptides which infect the brain. We found that, in many cases, families of similar inserted peptides yielded similar transduction rates (**Fig. 5c**). This effect was not limited to a particular capsid or insertion site, insofar as inserted peptides were often functional across all tested capsids and insertion sites (**Fig. 5c**). The observation that inserted peptides yielded consistent phenotypes across multiple capsids suggests that re-targeting is directly due to a peptide mediated mechanism. This result also highlights the power of peptide tiling library designs, insofar as having multiple overlapping peptides with similar sequences can function as internal controls, imparting confidence in the identification of particular AAV variant hits.

We then specifically interrogated two brain targeting variants (AAV5.APOA1 and AAV9.APOA1) which contained the same inserted apolipoprotein-A 1 (APOA1) derived peptide. Both variants were confirmed to package into functional capsids by electron microscopy (**Supplementary Fig. 3**). Across the entire APOA1 protein, functional APOA1-derived peptides were primarily from a tandem repeat region at the C-terminus of the protein (**Supplementary Fig. 3**). Highlighting the peptide-specific effect of altered AAV tropism, we observed strong detargeting away from the liver at less than 2% of AAV9 levels for both variants, with unperturbed brain transduction relative to wild-type AAV9 (**Fig. 4c**). This was especially remarkable for AAV5.APOA1 as this engineered variant was able to overcome the limited transduction of its parental AAV serotype across the blood-brain barrier. AAV5.APOA1’s improved specificity was subsequently shown to be strain independent as it retained both its brain targeting and liver detargeting functionality in BALB/c mice **(Supplementary Fig. 2b)**.

### Engineering re-targeted immune orthogonal AAV vectors

To generate additional variants with clinically relevant features, we next sought to expand our AAV engineering approach to include novel capsids beyond AAV5 and AAV9. In particular, we explored the potential of engineering tropism of highly divergent AAV orthologs. This was motivated by the fact that one of the biggest challenges with AAV therapy is the associated immune responses: either, pre-existing immunity that currently excludes a significant fraction of the human population from being eligible for any AAV therapy (*69*); or induced immunity upon AAV injection which prevents subsequent AAV re-dosing. This makes it extremely difficult to create effective therapies since it only allows a single opportunity to treat a patient, thus often mandating extremely high titers for a one-time therapy which in turn can lead to dangerous levels of toxicity. Recently (*70*), we proposed the concept of immune orthogonality to address this challenge. Specifically, we suggested that an orthologue, given sufficient sequence divergence, may not cross-react with the immune response generated by exposure to other orthologues, thereby potentially allowing re-dosing that avoids neutralization by existing antibodies or clearance of treated cells by activated cytotoxic T cells. While in that earlier study we focused on commonly used AAV orthologs, we decided here to explore highly divergent natural orthologs from the full spectrum of mammalian species, including those which have not yet been thoroughly tested for use as *in vivo* vectors. Towards this we established an integrated computational-cum-experimental pipeline to identify novel immune-orthogonal AAV serotypes. Specifically, first, basic local alignment search tool (BLAST) was used to identify 687 capsids with sequence homology to the AAV2 *cap* gene. This initial list was then filtered to exclude truncated genomes, redundant samples, human and non-mammalian serotypes, as well as close orthologs, resulting in 23 AAV capsid sequences for experimental investigation **(Fig. 6a, Methods).** Analyzing the evolutionary distance of these potential AAV orthologs, we confirmed that they exhibit high dissimilarity from all wild-type AAV capsids (except AAV5 which we had previously shown to be immune-orthogonal (*70*)), with many exhibiting less than 60% sequence similarity to AAV2 (**Fig. 6b**). To investigate if these computationally predicted AAV orthologs were functional, we first checked their ability to package into capsids. Comparing titers to AAV5, we identified 11 of the 23 AAV capsids with packaging titers within 10-fold of AAV5 (**Fig. 6c**). Next, to evaluate *in vivo* transduction potential, these 11 AAV capsids carrying a mCherry transgene were administered individually to C57BL/6 mice. Given that most AAV capsids have at least a degree of liver transducing potential, we harvested the livers of these mice 3 weeks post infection. Of the 11 AAV capsids analyzed, 4 of them had detectable levels of mCherry in the liver with 2 of them generating 3-fold higher mCherry expression than AAV5 (**Fig 6c**).

**Figure 6.**
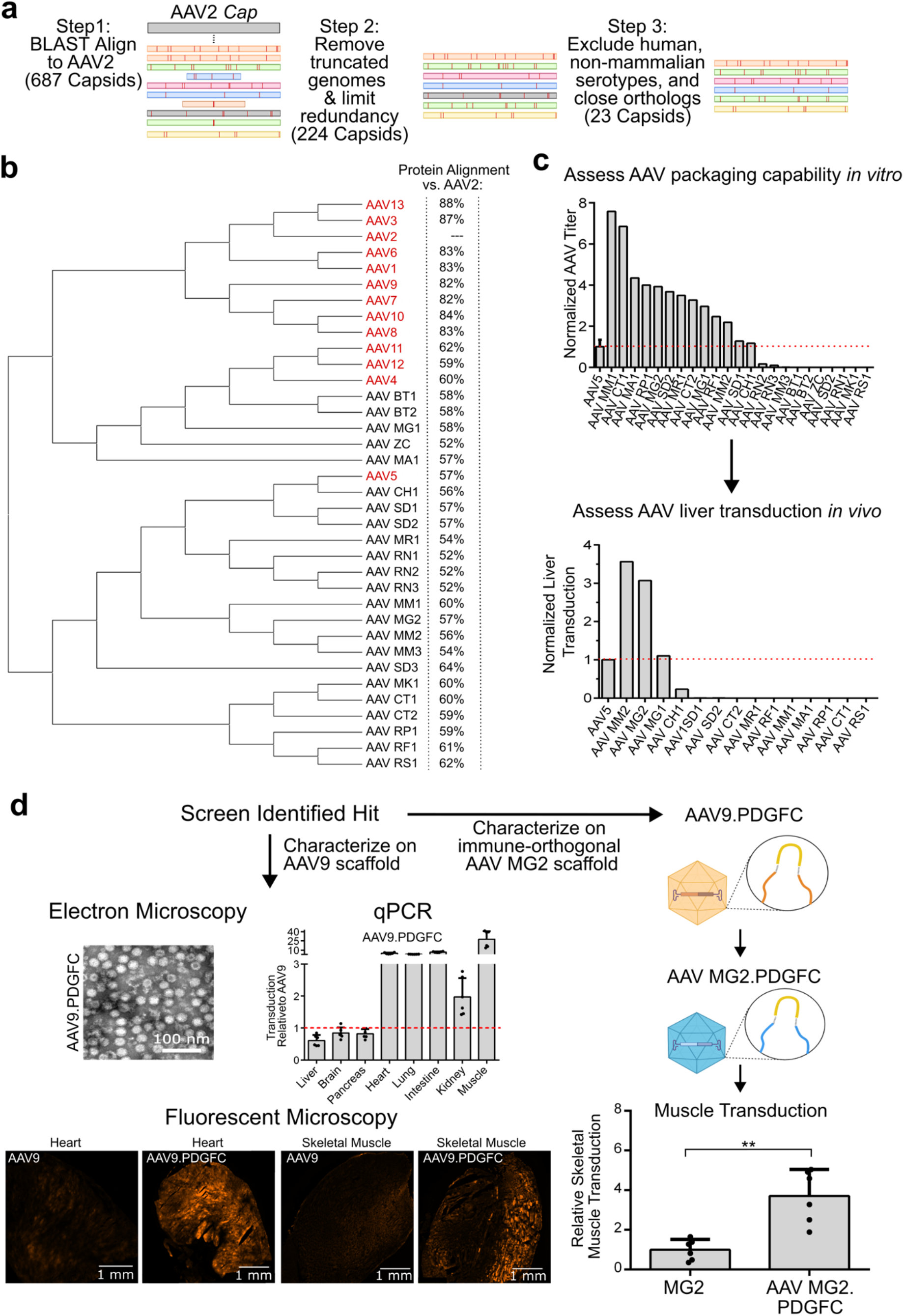
Novel immune-orthogonal AAV serotype mining, characterization and tropism reprogramming via displayed ligand peptides. **(a)** Schematic of the computational pipeline used to identify novel AAV serotypes for testing. Basic local alignment search tool (BLAST) was used to identify 687 initial capsids with sequence homology to the AAV2 *cap* gene. This initial list was then filtered to exclude truncated genomes, redundant samples, human and non-mammalian serotypes, as well as close orthologs. The final resulting list contained 23 AAV capsid sequences for subsequent investigation. **(b)** Hierarchical clustering dendrogram of AAV capsid sequences. Shown in red are previously identified AAV serotypes currently in use. Novel AAV capsids identified and used for downstream testing are shown in black. **(c)** Two stage filtering of novel AAV capsids: AAV capsids were first assessed by measuring their ability to package, and then by their ability to transduce the liver (using an mCherry transgene) *in vivo.* All values shown relative to the orthologous wild-type AAV5. **(d)** Characterizing muscle-targeting AAV variant in the AAV9 scaffold and assessing the feasibility of peptide transfer to an immune-orthogonal AAV for re-targeting tropism: Shown on the left are the full characterization experiments for the variant AAV9.PDGFC. Bar plot shows the RT-qPCR quantification (n=2) of mCherry transgene expression via individual validation, normalized to that of AAV9. Also shown are electron micrographs for variant capsids, as well as fluorescent microscopy of mCherry protein expression levels in the heart and muscle. The PDGFC peptide from AAV9.PDGFC was inserted onto loop1 of AAV MG2 to yield AAV MG2.PDGFC **(Methods)**. We injected AAV MG2 and AAV MG2.PDGFC into C57BL/6 mice, and quantified muscle transduction via RT-qPCR after three weeks. Bar plots show muscle transduction relative to wild-type AAV MG2. Statistical significance between groups was calculated via a t-test (*p<0.05, **p<0.01, ***p<0.001, ****p<0.0001).

Having identified 4 functional AAV capsids, we next characterized their immune-orthogonality by injecting C57BL/6 mice with each of these and collecting serum at day 0 and day 21 timepoints **(Supplementary Fig. 4a, Methods).** AAV8 was selected for comparison in this study due to its propensity for transducing the liver and its similar serological profile to other wild-type serotypes (*71*). Performing an ELISA at the 3-week timepoint confirmed that antibodies developed against the 4 studied immune-orthogonal AAV capsids indeed exhibited no immune cross-reactivity with AAV8 **(Supplementary Fig. 4a)**. This result highlights the utility in combining computational and experimental methods to identify and characterize novel AAV serotypes which could be repurposed to enable gene therapy redosing (*70*).

Importantly, next we assessed whether the tropism of our mined immune-orthogonal AAV orthologs could be re-engineered through insertion of one of our screen identified hit peptides. As we had observed that our muscle targeting AAV variants had broader tissue-tropism than the brain and lung targeting variants, insofar as they also readily infected the heart, lung, intestine, and spleen at levels comparable to, or exceeding, AAV9, we hypothesized that grafting the peptide from one of these variants onto an AAV ortholog would result in enhanced muscle targeting with minimal effects on parental AAV infectivity. To assess this, we selected AAV9.PDGFC as a candidate variant, and confirmed its ability to form functional capsids **(Fig. 6d, left)** and enable efficient targeting of muscular tissue (with >10 fold expression above AAV9 as quantified by RT-qPCR) **(Fig. 6d, left)**. The skeletal muscle and cardiac muscle (heart) transduction enhancement relative to AAV9 was also confirmed by protein level visualization **(Fig. 6d, bottom left)**. We then grafted this 20-mer PDGFC derived peptide onto one of the surface loops of the immune orthogonal AAV MG2 **(Fig. 6d, right, Methods)**, and notably observed similar enhanced muscle transduction of the engineered variant above its parental AAV **(Fig. 6d, right).** Taken together, this result highlights the fidelity of our ligand tiling display approach and lends further credence to the exciting possibility that displayed peptides identified in our screening approach can be utilized to re-engineer a diverse clade of AAV serotypes via peptide transfer.

## DISCUSSION

Rational screening strategies have immense potential to expand the molecular tools available for clinical gene therapy applications (*72*). While AAV engineering efforts have been conducted for over a decade (*28, 73*), advances in DNA synthesis have enabled us to create here a data-driven library of AAV variants leveraging existing functional biomolecules from nature (**Fig. 1**). Using natural biomolecules as a defined source of inserted peptides has multiple benefits over random hexamers (and similar methods). First, natural biomolecules have been pre-filtered for biological functionality by millennia of evolutionary selection pressure. Second, a defined library allows for robust quantification of the fitness of each AAV variant, enabling facile stratification of AAV variants by infectivity across organs of interest. While we primarily applied this methodology to engineer AAVs, mining nature for functional biomolecules has applications in a wide range of protein engineering challenges, such as engineering orthogonal viral (including lentiviruses) and non-viral (including lipid nanoparticles (*74*)) vectors or identifying biologic inhibitors of critical protein/protein interactions, as we have shown recently (*37*).

Additionally, we believe that the rational ligand tiling approach that we developed herein could lead to important insights into basic AAV-receptor binding and transduction more generally. Previous efforts have either utilized large-scale genetic screens (*75*) or unique observations about a particular AAV variant’s transduction behavior (*76, 77*). As the AAV variants that we have described display peptides derived from natural receptor-interacting ligands, we believe that our AAV variants may maintain some of this binding capacity which partially explains their altered transduction profiles. To this end, we have shown that one of our discovered variants (AAV9.DKK1) exhibits deprecated transduction when the cognate receptor for the ligand from which the peptide was derived is knocked out (**Fig. 5a,b**). Furthermore, due to the systematic tiling nature of our peptide library, we were able to generate full-length transduction maps across the entire residue space of all ligands utilized in this study **(Fig. 5a, Supplementary Fig. 3)**. This not only increases confidence in our screening hits, in that homologous peptides behave similarly, but could also provide insight into critical structural domains mediating virus-receptor interactions (*78, 79*) and more broadly, provide a platform for further expanding the known protein interactome (*80, 81*) especially when our dataset is integrated with the cell surfaceome (*82, 83*).

Additionally, in recent years, developing predictive models of AAV infectivity has garnered significant interest from multiple research groups. The application of machine learning to AAV engineering parallels major advances in machine learning across multiple areas of protein science such as structure prediction (*84–86*), enzyme activity forecasting (*87, 88*), and antibody binding optimization (*64*). While deep learning and similar blackbox methodologies have rapidly become mature technologies, applying these methodologies to AAV engineering is still severely limited by the lack of available training data. Our AAV screening data is an ideal training dataset for several reasons: (1) our experimental design features a large, defined library of variants (**Fig. 1**), meaning that every variant has a reliable quantification of packaging and infectivity, (2) we screened each variant across a panel of 9 major organs to map the infectivity across diverse tissue types, (3) our library was inserted across 2 AAV serotypes and 2 surface loops, providing important functional information on the capsid context in which the peptide resides, (4) we have rigorously, individually validated a large cohort of variants to demonstrate our screening data is trustworthy (**Fig. 4**); and (5) to illustrate the utility of our dataset as training data, we demonstrated the packaging efficiency of AAV variants could be accurately predicted from the biophysical characteristics of the inserted peptides (**Fig. 2d**), and also peptide amino acid sequence is directly predictive of tissue-tropism across multiple capsids and insertion sites (**Fig. 3c**). As such, we anticipate our screening data will have great utility for the machine-learning and computational biology community.

While the variants we identified via our pooled screen have tissue transduction exceeding AAV9 in many organs (**Fig 5a**, **Fig. 6d**) and several variants exhibit drastic liver de-targeting **(Supplementary Fig. 2a)**, further engineering could be performed to enhance potency and specificity. This may be relevant in particular for certain peptides derived from ligands that engage receptors expressed on multiple cell types or which have promiscuous binding activity. In our validation experiments, we used a standard promoter (CMV) to drive expression of the mCherry transgene. Alternatively, tissue-specific promoters could be used to increase the specificity and magnitude of transgene expression in the organ of interest (*18, 89, 90*). Furthermore, the hit capsids identified here could be further engineered for increased activity. Existing hits could serve as a scaffold for further rounds of targeted mutagenesis and screening, or peptides could be inserted on both loop1 and loop2 of the AAV capsid to increase the valency of the displayed ligands (*91*). Additionally, recently developed direct chemical engineering (*31*), or peptide display strategies (*92*), and/or alternative peptide linkers (*93, 94*) could be utilized to enhance peptide-mediated transduction. Also, scRNAseq could be used to screen hit variants towards more specific cell-types within the organ of interest (*95, 96*).

Finally, while our core screening strategy addresses the gene therapy hurdle of organ targeting, the issue of pre-existing AAV immunity can limit clinical translation (*69*). Therefore, we computationally identified and fully characterized a set of 4 immune-orthogonal AAV capsids **(Fig. 6a-c, Supplementary Fig. 4a),** and we show that insertion of a screen identified re-targeting peptide can alter the *in vivo* tropism of one of these identified AAV capsids (**Fig. 6d**). This not only increases the number of known serotypes with interesting immunological properties to be investigated alongside the current AAV set, but also lends credence that the peptide insertions identified through our screening methodology can be grafted onto diverse biological scaffolds to achieve unique targeting across broad delivery contexts.

Taken together, we have presented a massive functional screen of engineered AAV variants, spanning over one million total variants derived from two capsids and multiple sites of insertional mutagenesis. Using this screening data, we individually validated 21 AAV variants, identifying AAV capsids with increased organ transduction across multiple organs (heart, muscle, and lung for AAV9.PDGFC, **Fig. 6d**), as well as highly specific AAV capsids (AAV5.APOA1, AAV9.DKK1, **Supplementary Fig. 3**, **Fig. 5a**). Improved broad targeting AAV variants have massive potential for genetic diseases such as hemophilia A, where total factor VIII expression levels are most critical. At the same time, highly specific AAV capsids such as AAV5.APOA1 (which has less than 1% the liver infectivity of WT AAV9) would have great utility for neurodegenerative disorders, where maximizing transgene expression in the brain is essential. In addition to the novel variants identified herein, the bulk screening data itself is high value. Given the scale, reliability, and translational relevance of our screening dataset, we anticipate it will serve as a foundation for future computational engineering of designer AAV capsids.

## ACKNOWLEDGEMENTS

This work was generously supported by UCSD Institutional Funds, a Department of Defense Grant (W81XWH-22-1-0401), and NIH grants (OT2OD032742, R01GM123313). The authors would like to thank the University of California, San Diego (UCSD) Cellular and Molecular Medicine Electron Microscopy Core (UCSD-CMM-EM Core) for equipment access and technical assistance. The UCSD-CMM-EM Core is supported in part by the National Institutes of Health (NIH) Award number S10OD023527. The authors would also like to thank the microscopy core in the UCSD neurosciences department which is supported by a NIH grant (NINDS P30NS047101), the La Jolla Institute Histology Core facility for their expert help with tissue preparation and cryosectioning, and the GT3 Core Facility of the Salk Institute which is supported with funding from NIH-NCI CCSG: P30 014195, a NINDS R24 Core Grant, and funding from NEI.

---

**Supplementary Figure 1:**
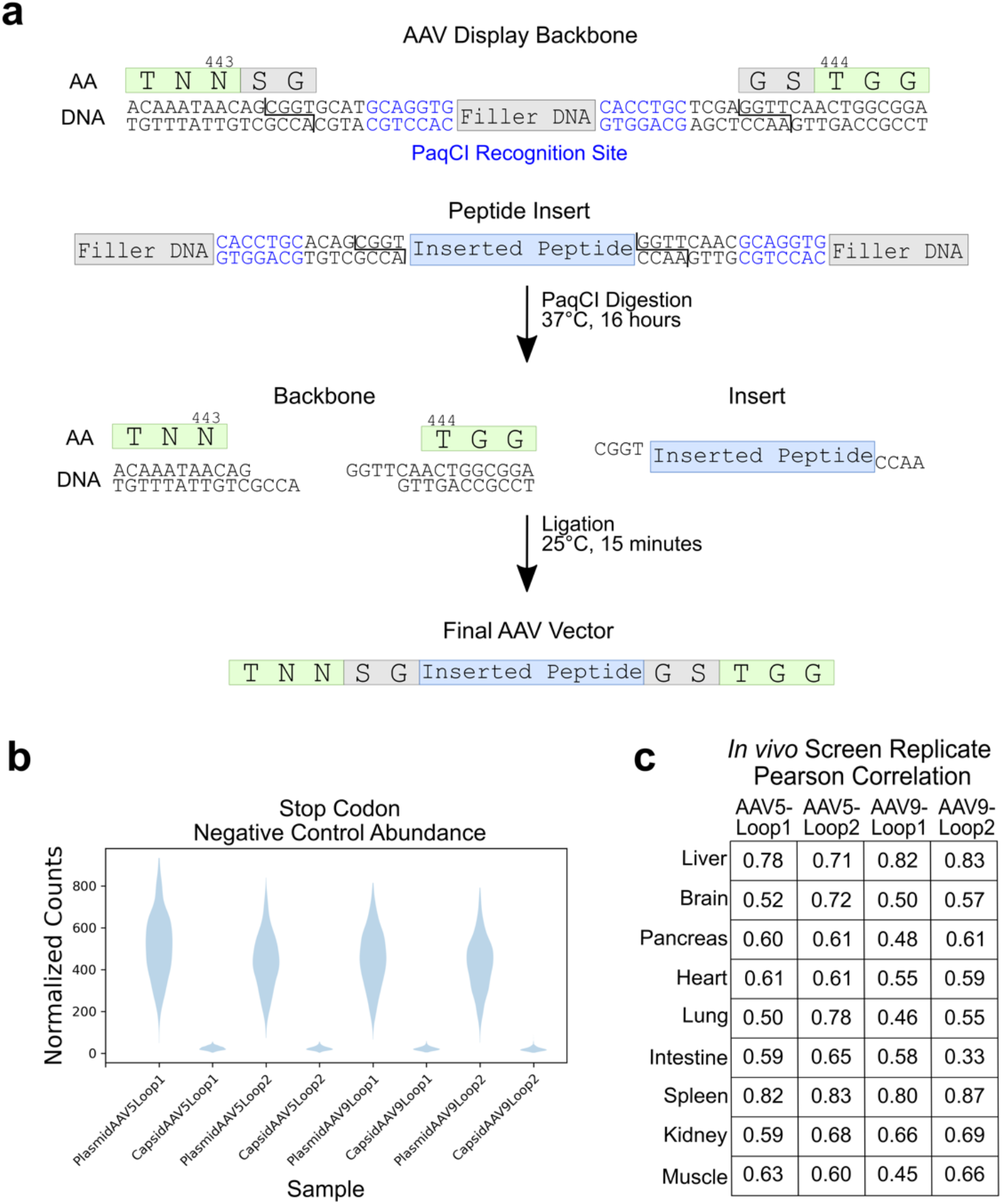
AAV peptide display library construction scheme, and library screening quality controls. **(a)** Schematic illustrating the detailed cloning procedure for inserting the peptide libraries derived from an oligonucleotide pool into the respective AAV capsid backbone. The AAV backbone was first modified at the insertion location to contain flanking glycine-serine linkers with type IIS PaqCI recognition sites. The synthesized peptide libraries were produced with PaqCI recognition sites on the 5’ and 3’ ends. Both the backbone and the peptide insertion library were digested overnight at 37℃. These were then ligated together to produce the final engineered AAV library **(Methods)**. Shown is the AAV5-Loop1 backbone for illustrative purposes. **(b)** Normalized abundance for negative control peptides containing stop codons, in both the plasmid libraries and recombinantly produced AAV capsid libraries for all four capsid-insertion site combinations. **(c)** Replicate Pearson correlation for the log_2_ fold change (log_2_FC) values (organ vs. capsid) for each library and organ.

**Supplementary Figure 2:**
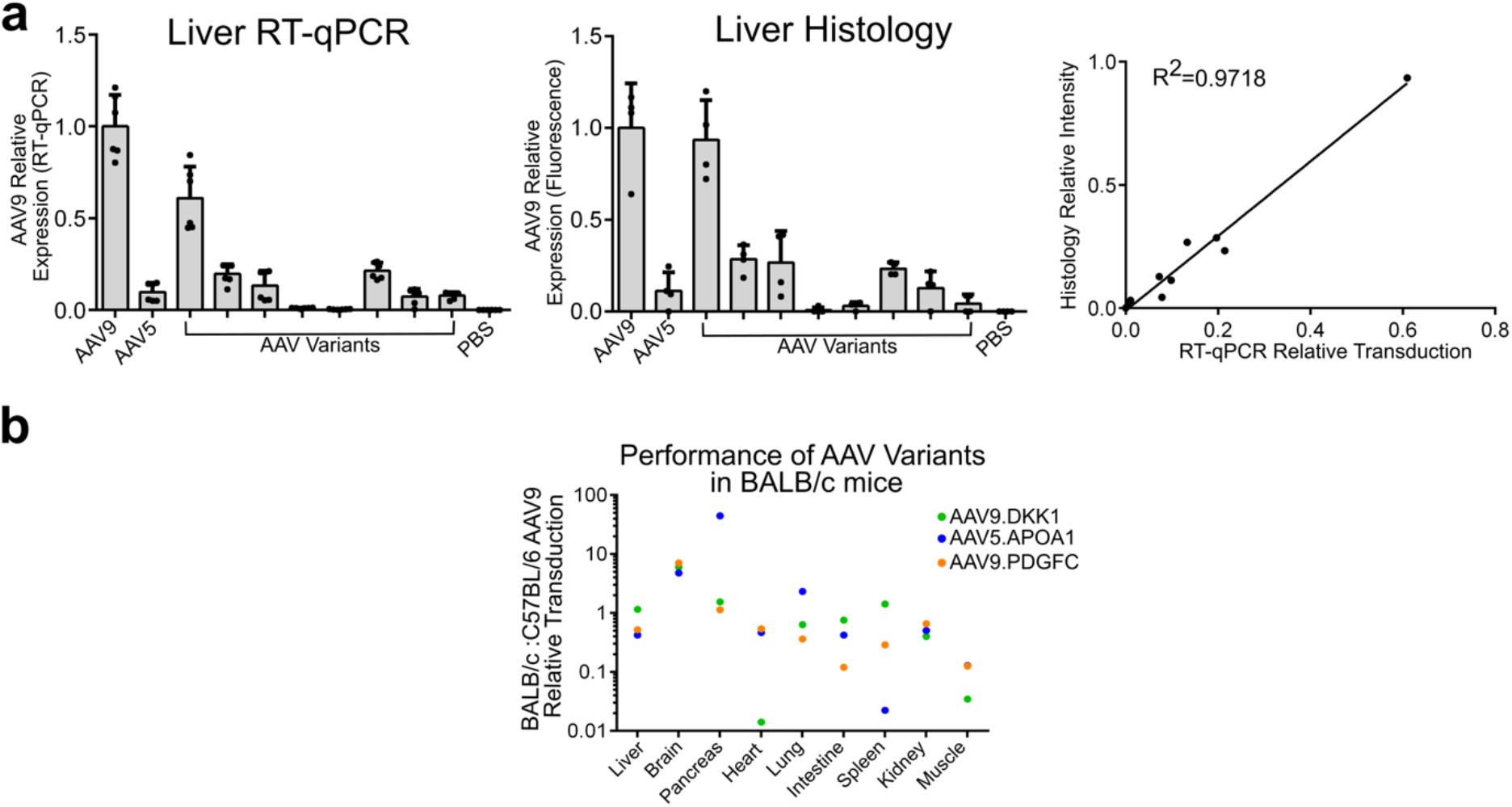
*In vivo* validations of AAV variants assayed via RNA and protein measurements, and across mice strains. **(a)** Comparison between liver RT-qPCR quantification of mCherry delivery, versus protein level quantification via fluorescent microscopy **(Methods)**. **(b)** Dot plot showing the concordance of three individually validated AAV variants assessed in BALB/c mice. The relative organ transduction levels for all variants were quantified relative to AAV9 in both strains and then a BALB/c to C57BL/6 relative transduction ratio was calculated and plotted.

**Supplementary Figure 3:**
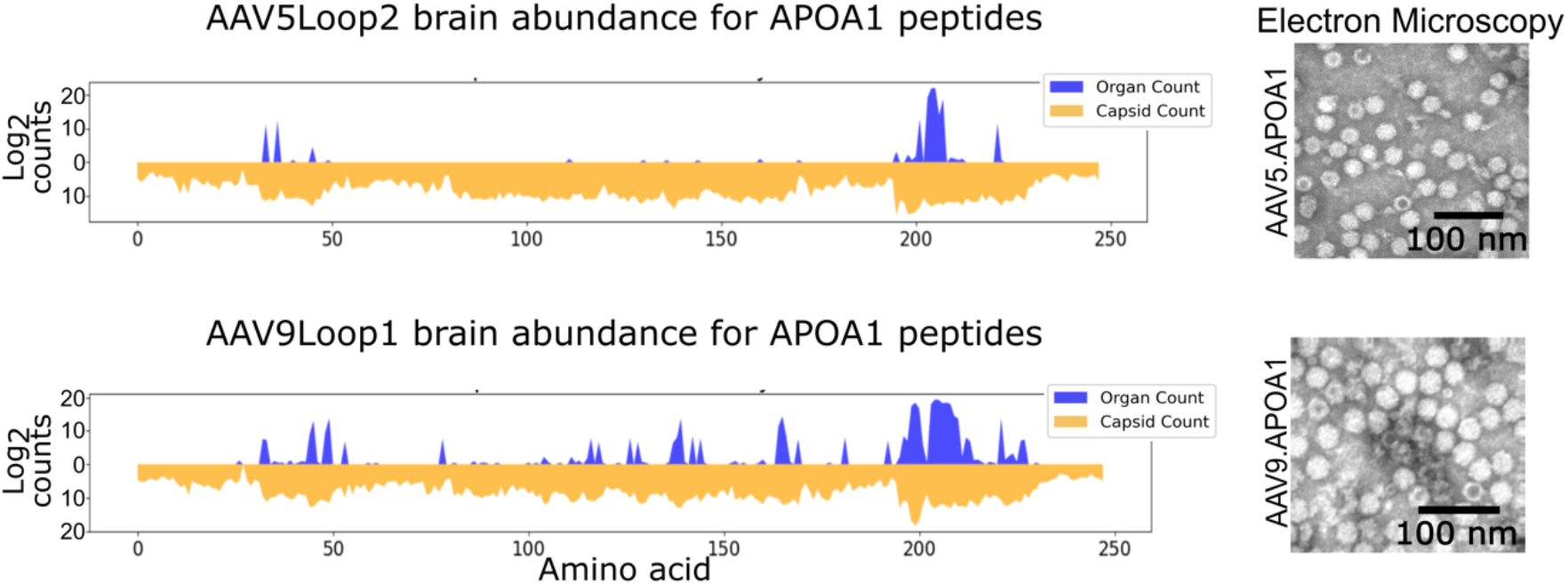
*In vivo* screen identified brain specific AAV.APOA1 variants. Brain and capsid counts for all AAV5Loop2 and AAV9Loop1 variants with inserted APOA1 derived peptides. Shown in blue are the brain counts, and shown in orange are the capsid counts. Also shown are electron micrographs for AAV5.APOA1 and AAV9.APOA1.

**Supplementary Figure 4:**
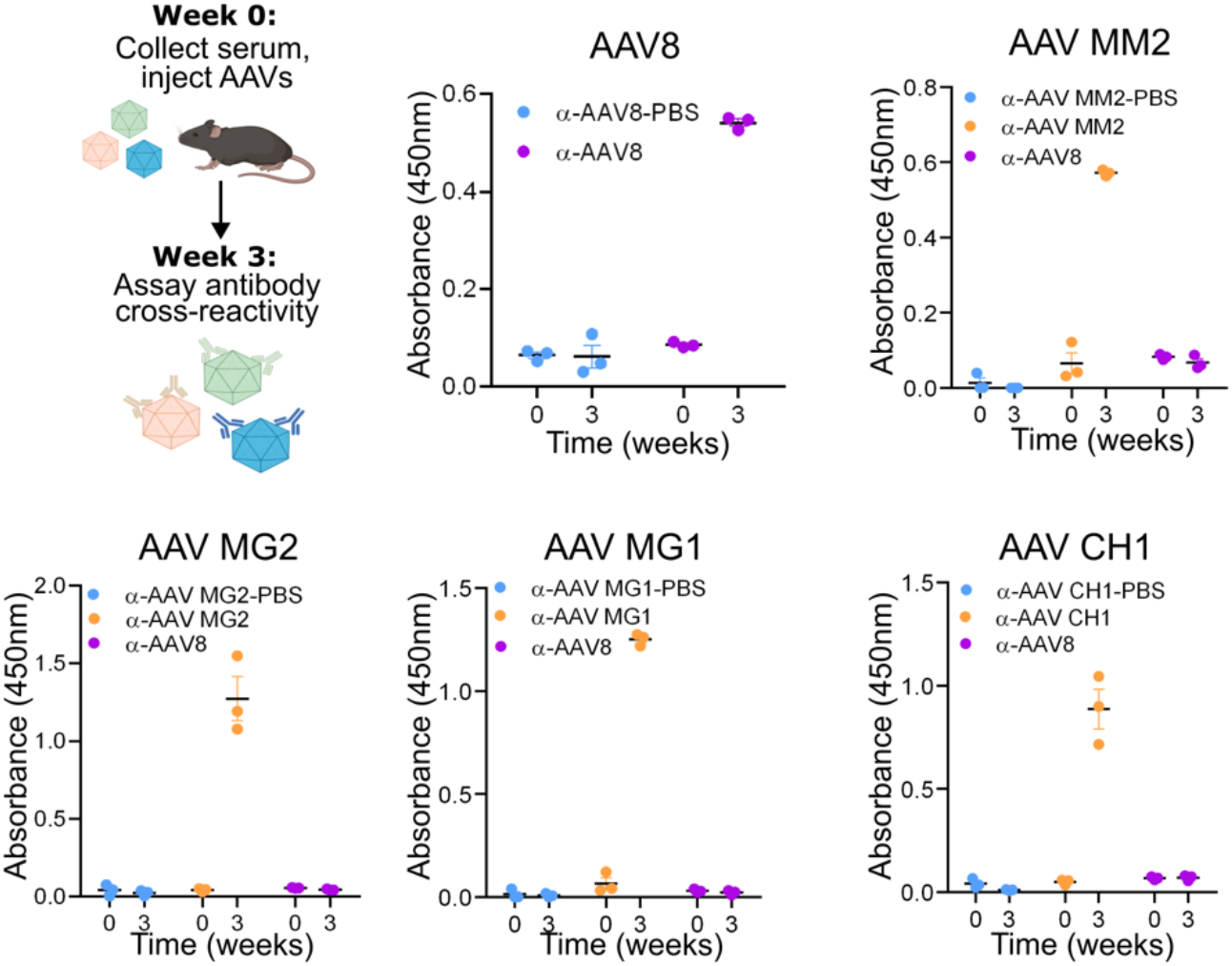
Computationally identified AAVs demonstrate robust immune-orthogonality when compared to AAV8. The four novel AAV capsids which could infect the liver (AAV MM2, AAV MG2, AAV MG1 and AAV CH1), were tested for immune cross-reactivity with AAV8. Mice were immunized with the indicated AAV, and then 3 weeks post-injection tested for antibody cross reactivity via an ELISA (**Methods**).

## MATERIALS AND METHODS

### Mice

All animal care and experimental methods were performed in accordance with the University of California Institutional Animal Care and Use Committee. 6-8 week old male C57Bl/6J (JAX, #000664) and Balb/cJ (JAX, #000651) mice were purchased from the Jackson Laboratories and systemic injections were administered retro-orbitally with either AAV or PBS.

### Cell lines and culture conditions

All cells were cultured in a 37°C 5% CO2 humidified incubator. HEK293T cells were cultured in DMEM medium supplemented with 10% FBS, GlutaMAX (1x) (GIBCO), and Penicillin-Streptomycin (100 U/mL) (GIBCO).

### Visualizing AAV capsid structures

To obtain structural representations of AAV capsids, AAV5 (*65*) and AAV9 (*66*) structure files were downloaded from the Protein Data Bank. These were visualized using the PyMOL Molecular Graphics System, Version 2.0 Schrödinger, LLC with the *ramp_new* function utilized for coloring the surface capsid representation.

### Design of displayed peptide library

Each AAV library consisted of 275,298 peptides, derived from 6,465 proteins. These protein sources were mined from a variety of protein families, including all protein ligands cataloged in the Guide to Pharmacology database (*48*), toxins, nuclear localization signals (NLS), viral receptor binding domains, albumin and Fc binding domains, transmembrane domains, histones, granzymes, and predicted cell penetrating motifs. In addition to peptides coding for functional biomolecules, we also included 444 control peptides coding for FLAG-tags with premature stop codons. For all human proteins, the cDNA coding for each protein was fragmented *in silico* to generate DNA coding for all possible 20mer peptides. For viral proteins, cell penetrating motifs, and FLAG stop codon controls, the protein sequence was back-translated to DNA using the most abundant human codon for each amino acid.

### Oligonucleotide array synthesis, amplification and cloning

Oligonucleotide libraries were synthesized by GenScript as three 91,766 element pools. Each oligonucleotide library was amplified using KAPA Hifi Hotstart Readymix and the manufacturer recommended cycling conditions with an annealing temperature of 60 °C and an extension time of 30 seconds. The number of PCR cycles was optimized to avoid over-amplification of the peptide libraries. After amplifying each oligonucleotide pool and confirming amplicon size on an agarose gel, the amplified sub-libraries were pooled to yield the total 275,298 element peptide library. These pools were cloned into loop 1 or loop 2 sites to yield pAAV5L1_Screen, pAAV5L2_Screen, pAAV9L1_Screen, and pAAV9L2_Screen. The AAV *rep* and *cap* were flanked by AAV inverted terminal repeat (ITR) sequences to facilitate packaging of *cap* genes into a recombinant AAV particle.

### Recombinant AAV production

Utilizing the library plasmid pools described above (AAV5-Loop1, AAV5-Loop2, AAV9-Loop1, and AAV9-Loop2), each AAV capsid library was produced by transfecting HEK293T cells in 40 15 cm dishes with the plasmid library pool (diluted 1:100 with pUC19 filler DNA to prevent capsid cross-packaging) and an adenoviral helper plasmid (pHelper) (*97*). Titers were determined via qPCR using the iTaq Universal SYBR green supermix and primers binding to the AAV ITR region. To prepare the capsid particles as templates for qPCR, 2 μL of virus was added to 50 μL of alkaline digestion buffer (25mM NaOH, 0.2 mM EDTA) and boiled for 8 minutes. Following this, 50 μL of neutralization buffer (40mM Tris-HCl, .05% Tween-20, pH 5) was added to each sample.

### In vivo evaluation of AAV display libraries

Each AAV capsid library was retro-orbitally administered to mice in duplicate at a dose of 2E12 vg/mouse for the AAV9-based libraries or 1E12 vg/mouse for the AAV5-based libraries. Two weeks after injection, the heart, lung, liver, intestine, spleen, pancreas, kidneys, brain, and gastrocnemius muscle were harvested and placed in RNAlater storage solution. Total DNA was extracted from all mouse tissues using TRIzol reagent and the TNES-6U back extraction method (*98*). The resulting precipitated DNA was centrifuged for 15 minutes at 18,000G, and the supernatant discarded. The DNA pellet was then washed three times with 70% ethanol, and finally resuspended in 300 μL of EB after allowing the pellet to air dry.

### Preparation of plasmid and capsid DNA for next generation sequencing

To sequence the plasmid libraries (AAV5/9 and loop1/2 peptide insertions), 50 ng of plasmid was used as template for a 50 μL KAPA Hifi Hotstart Readymix PCR reaction with melting temperature of 60°C and an extension time of 30 seconds. The primers were designed to amplify the peptide coding region from each sub-library. The number of cycles was optimized to avoid overamplification. The PCR reactions were purified using a QIAquick PCR Purification Kit according to the manufacturer’s protocol. Following this, 50 ng of the PCR amplicon was used as template for a secondary 50 μL KAPA Hifi Hotstart Readymix PCR reaction to add illumina compatible adapters and indices (NEBNext Cat# E7600S). The PCR reaction was performed with a melting temperature of 60°C, an extension time of 30 seconds. To sequence the capsid libraries, a similar protocol was performed, with a modified template amount in the step-1 PCR. To prepare the capsid particles as templates for PCR, 2 μL of virus was added to 50 μL of alkaline digestion buffer (25mM NaOH, 0.2 mM EDTA) and boiled for 8 minutes. Following this, 50 μL of neutralization buffer (40mM Tris-HCl, .05% Tween-20, pH 5) was added to each sample. 1 μL of this digested capsid mix was then used as a template for a 50 μL PCR reaction. For each sample, the number of cycles was optimized to avoid overamplification, and a secondary PCR was subsequently performed to add illumina compatible adapters and indices. After generating illumina compatible libraries, the plasmid and capsid samples were sequenced on a NovaSeq 6000 with an S4 flowcell generating 100bp paired end reads.

### Preparation of tissue DNA for next generation sequencing

To sequence the AAV *cap* genes from each tissue for the pooled screen, as with the plasmid/capsid libraries, a two step PCR based library prep method was used. For each organ and replicate, a 300 μL PCR reaction was performed with 120 uL of genomic DNA used as a template. For each tissue, the number of cycles was optimized via an initial qPCR to avoid overamplification of the library. All other parameters such as primers, and melting temperatures were identical to the PCRs for the plasmid libraries. Following this initial PCR, a secondary PCR was performed as above to add illumina compatible adapters and indices. The libraries were then sequenced on a NovaSeq 600 with an S4 flowcell generating 100bp paired end reads.

### In vivo validation of AAV variants

Either saline or the AAV-variant-mCherry, AAV9-mCherry, or AAV5-mCherry capsids were systematically administered to mice in duplicate at a dose of 5E11 vg/mouse. Three weeks after injection, the lungs were inflated with a PBS/OCT solution and the lungs, heart, liver, intestine, spleen, pancreas, kidneys, brain, and gastrocnemius muscle were harvested. Each organ was split with one portion placed in RNAlater and the other embedded in OCT blocks and flash frozen in a dry-ice/ethanol slurry. Total RNA was then isolated from all mouse tissues using TRIzol reagent and RNA Isolation kits with on-column DNase treatment (Zymo Cat# R2072). cDNA synthesis was performed with random primers from the Protoscript cDNA synthesis kit (NEB Cat#E6560S). Transgene expression was then quantified via qPCR using the iTaq Universal SYBR green supermix and primers binding to the mCherry transcript. mCherry transgene expression was normalized to that of an internal GAPDH control, using GAPDH specific primers. For histological examination, OCT frozen blocks were cryosectioned at approximately 10 μm thickness and tissue slides were then imaged on an Olympus SlideScanner S200. Exposure times between 5-1000 ms were used, with identical exposure times used for all samples of a given tissue type. mCherry expression from histological sections was then quantified using the Olympus OlyVIA software to calculate the mean pixel intensity across the entire organ section.

### Quantifying AAV variant abundance from NGS data

Starting with FASTQ sequencing files, the MAGeCK (*99*) ‘count’ function was used to generate count matrices describing AAV abundance in each sample (plasmids/capsids/tissues). Following this, the count matrices were normalized (via multiplication with a constant size-factor) for each sample to account for non-identical read depth. The sequencing counts were then transformed by taking the log base 2 of the raw counts, after addition of a pseudocount. Variants with no counts across all of the experimental samples were excluded from analysis.

### Biophysical analysis of AAV capsids

The biophysical characteristics of the inserted peptides was calculated using the “ProteinAnalysis” module within the Biopython Python package (*100*). A variant was considered a successful packager if it had higher abundance in the capsid particles compared to the plasmid pool. Support vector machine training and visualization was accomplished via the “svm” module within the sklearn Python package (*101*). UMAP projection of peptide biophysical characteristics was accomplished via the “plot” functionality within the UMAP Python package (*102*). All default parameters were used for the visualization. Boxplots and hexbin plots were generated using the matplotlib and seaborn Python packages (*103*).

### Identifying significantly enriched variants in each tissue

To identify variants which successfully transduce each tissue, for each variant a one sample T-test was applied, comparing the abundance in the capsid particles to the abundance in the tissue. Resulting p-values were adjusted for multiple hypothesis testing via the Benjamini-Hochberg procedure (*104*). A variant was considered a significant transducer of an organ if it had an FDR adjusted p-value < .05, and a Log_2_FC > 1 in both replicates. When choosing variants for validation experiments, we prioritized variants which had inserted peptides which were identified as hits in multiple capsid/loop contexts, and variants for which we identified similar inserted peptides infecting the same organ.

### Visualizing tissue transduction from pooled screen

Heatmaps for visualizing AAV transduction were generated using the ‘clustermap’ function within the seaborn Python package (*103*). Rows and columns were ordered via the scipy ‘optimal_leaf_ordering’ function to minimize the euclidean distance between adjacent leaves of the dendrogram. UMAP projections visualizing AAV tissue specificity were generated by embedding the tissue level log_2_ fold change into two dimensions via the “plot” functionality within the UMAP Python package (*102*). All default parameters were used for generating the embedding. The variants were colored by the organ in which they had the max log_2_ fold change.

### Assessing accuracy of predicted AAV variant tropism

For each variant which was individually validated, we assessed the accuracy of both positive and negative predictions of tissue infectivity. For variants predicted to target a specific organ, we considered a prediction accurate if the individual validations showed greater than 50% of wild-type AAV9 infectivity in that organ. For variants predicted not to target a specific organ, we considered a prediction accurate if the individual validations showed less than 50% of wild-type AAV9 activity.

### Peptide Distance Projections

To calculate the Levenshtein distance between inserted peptides, the “levenshtein” function from the Python package “rapidfuzz” was used with default parameters(*105*). After building the pairwise distance matrix between all significantly enriched peptides, the matrix was projected into two dimensions via UMAP with metric=“precomputed”, n_neighbors=1500, and min_dist=.1. Clusters of peptides with similar functions were then hand annotated onto the resulting plot.

### Convolutional Neural Networks

To train convolutional neural networks (CNNs) to predict the tissue specificity of AAVs, we first converted the AA sequences of the inserted peptides to a one-hot encoding via the “get_dummies” function from the pandas Python package(*106*). Among the significantly enriched variants, a variant was considered a transducer of a given organ if the log_2_FC relative to the capsid in both replicates was greater than 0. The data was then randomly split into training (⅔) and validation (⅓) datasets. For each variant, the one-hot encoding was reshaped to a 20×20 matrix with rows indicating residue positions, and columns indicating the presence or absence of a particular amino acid. The model architecture was instantiated via a Keras sequential model(*107*). In brief, a convolutional layer (Conv1D) with 32 filters, a kernel size of 3 and “relu” activation was fed into a max pooling layer (MaxPool1D) with pool size of 2. These layers were followed with another set of convolutional and max pooling layers, this time with 64 filters in the convolutional layer. These layers were followed with a dense layer with units=20. Finally a dropout layer was added with the dropout rate=.5. A flattening layer and final dense layer (with sigmoid activation) was then used to output resulting class probabilities. A separate independent model was trained for each organ. When training the models, the classes (infective versus non-infective variants) were weighted proportionally to the inverse of the number of class examples. When calling ‘model.fit()’ to train the CNN, a dictionary describing the class weights was passed via the ‘class_weight’ parameter. Model performance was evaluated via accuracy, area under the receiver operator characteristic curve (AUROC), F1-score, and Matthews Correlation Coefficient (MCC). Metrics were calculated via builtin Keras functions, and plotted via matplotlib. To assess how model accuracy changes as a function of the edit distance from the training data, the levenshtein distances between peptides in the testing and training datasets were calculated using the ‘rapidfuzz’ python package as above.

### Transmission electron microscopy (TEM)

To obtain transmission electron microscopy images of select AAV variants, 20µL of the AAV solution was applied to formvar/carbon coated EM grids. Upon washing 3 times with water, the sample-containing grid was then negative-stained for 1 minute using a solution of 2 % of Uranyl acetate in water and blot-dried. The EM grids with each AAV variant were then imaged at 68,000X magnification using the FEI Tecnai Spirit G2 BioTWIN transmission electron microscope operated at 80keV.

### Mechanism knockout experiments

sgRNA sequences targeting LRP6 or non-targeting controls were identified using CRISPick (*108, 109*) and cloned into the lentiCRISPR v2 plasmid backbone (*110*). Lentivirus was then produced as described in (*37*). Briefly, HEK293T cells were seeded at ∼40% confluency the day before transfection. The day of transfection, Optimem serum reduced media was mixed with Lipofectamine 2000 (Thermo Fisher), 3 µg of pMD2.G plasmid, 12 µg of pCMV deltaR8.2 plasmid, and 9 µg of the respective lentiCRISPRv2 plasmid and added drop-wise onto the HEK293T cells following a 30 minute incubation period. 48 hours after transfection, the media was collected and replaced with fresh DMEM with 10% FBS. At 72 hours post-transfection, the supernatant containing viral particles were collected again, pooled, and concentrated to 1mL using Amicon-15 centrifugal filters with a 100,000 NMWL cutoff (EMD Millipore). For lentiviral transduction, HEK293T cells were seeded in a 12-well plate at ∼20% confluency the day before transduction. The day of transduction, lentiviral containing DMEM with 8µg/mL polybrene was added to the cells. The media was then replaced 24 hours later and then changed into puromycin (2µg/mL) containing DMEM 28 hours post-transfection. Post selection, once the cells reached confluency, they were passaged into a 24-well plate at ∼40% confluency. 24 hours later, DMEM containing either AAV9 (4×10^9^ viral genomes) or AAV9.DKK1 (1×10^9^ viral genomes) was added to the cells. The cells were then collected 24 hours later and flow cytometry was performed to quantify mCherry transgene expression.

### Identifying immune orthogonal AAV capsids

Potential AAV orthologs were curated by first identifying sequences which exhibited similarity to the AAV2 *cap* gene using the National Center for Biotechnology Information (NCBI) basic local alignment search tool (BLAST) (*111*). Next, incomplete, heavily truncated, and highly homologous sequences to one another were filtered out of the selection criteria. Viruses within the human AAV clade and those from non-mammalian hosts were then filtered out. Finally, sequences with high similarity to previously identified human serotypes were removed, resulting in the 23 potential AAV orthologues assessed.

### Immune orthogonal AAV production

To clone computationally identified AAV orthologs, the capsid sequences were codon-optimized and cloned downstream of the AAV2 *rep* gene using Gibson Assembly. Immune orthogonal AAV capsids were produced as described for the AAV variant validation capsids and AAV production titer was measured via qPCR with primers binding to the AAV ITR region. Any ortholog with a production titer within a power of 10 of AAV5 was considered to have successfully packaged.

### Immune orthogonal AAV in vivo transduction

Immune orthogonal AAV capsids with sufficient packaging titer were then injected retro-orbitally into C57BL/6 mice at a dose of 1×10^12^ viral genomes/mouse. Livers were harvested 3 weeks post-injection and total RNA was isolated as described above. cDNA was then generated and transgene expression was quantified via qPCR using the iTaq Universal SYBR green supermix and primers binding to the mCherry transcript. mCherry transgene expression was then normalized to GAPDH and the relative expression was compared to AAV5.

### Assessing immune cross-reactivity

The assay to assess immune antibody cross-reactivity of the identified immune orthogonal was performed as described in (*70*). Prior to injection, serum was collected via tail snip procedure and then the mice were injected with 1×10^12^ viral genomes/mouse of AAV or PBS in triplicate. 3 weeks later, serum was collected from each of the mice and the antibody cross-reactivity ELISA was performed. For this, 1×10^9^ viral genomes of AAV8, AAV MM2, AAV MG1, AAV MG2, or AAV CH1 were diluted in a 1x coating buffer and incubated overnight in each well of 96-well Nunc MaxiSorp plates. Plates were washed three times for 5 mins with 1x wash buffer (Bethyl) and blocked with 1x BSA blocking buffer (Bethyl) for 2 hours at room temperature. The wells were then washed again and serum samples were added at a 1:40 dilutions. Plates were incubated for 5 hours at 4℃ with shaking. Wells were 3x washed and 100 µL of HRP-labeled goat anti-mouse IgG1 (Bethyl; diluted 1:100,000 in 1% BSA) was added to each well. Secondary antibody was incubated for 1 hour at room temperature, wells were washed 3 times, and 100 µL of TMB substrate was added to each well. Optical density at 450 nm was measured using a microplate absorbance reader (BioRad iMark).

### Engineering immune orthogonal AAV variants

To engineer the immune orthogonal ‘AAV MG2’ capsid, we first aligned the AAV9 capsid amino acid sequence to the MG2 sequence using Clustal Omega (*112*). The PDGFC peptide coding sequence was then ligated into the ‘AAV MG2’ vector at the appropriate Loop 1 location with an identical protocol as was done for cloning peptides into AAV9. The engineered ‘AAV MG2.PDGFC’ vector capsid was then assayed *in vivo* identically to the above validation experiments using AAV9.

### Ethical Compliance

All experiments involving live vertebrates performed at UCSD were done in compliance with ethical regulations approved by the UCSD IACUC committee.

### Statistics

Unless otherwise noted, differences in means were calculated via an unpaired t test. When necessary, p values were adjusted for multiple hypothesis testing via the benjamini-hochberg procedure(*104*). For all Figures, unless otherwise noted: *p<0.05, **p<0.01, ***p<0.001, ****p<0.0001.

